# The Impacts of Historical Barriers on Floristic Patterns of Plant Groups with Different Dispersal Abilities in the Ryukyu Archipelago, Japan

**DOI:** 10.1101/007617

**Authors:** Koh Nakamura, Rempei Suwa, Tetsuo Denda, Masatsugu Yokota

## Abstract

The effects of historical barriers in biogeographical patterns are expected to persist differently depending on dispersal abilities of organisms. We tested two hypotheses that plant groups with different dispersal abilities display different floristic patterns, and that historical barriers can explain floristic differentiation patterns in plants with low dispersal ability but not in plants with higher dispersal ability, in the seed plant flora of the Ryukyu Archipelago. This area is biogeographically interesting because several similar floristic differentiation patterns have been proposed, all of which are primarily explained by two historical barriers, the Tokara Tectonic Strait (Tokara Gap) and the Kerama Gap, which arose during the formation of the islands. We calculated floristic dissimilarity distance among 26 islands based on data sets for three dispersal-ability classes. Clustering analyses based on the floristic dissimilarity distance generated similar floristic patterns regardless of dispersal-ability class. We propose that because the landscape resistance is so strong that migration of plants is severely restricted regardless of their dispersal abilities, the similar floristic differentiation patterns are generated. Multivariate regression analyses using Mantel’s randomization test indicated that floristic differentiations among islands were explained by the both effects of the historical barriers and geographic distance in all dispersal-ability classes. Significance of the historical barriers is not determined by the plant dispersal abilities but presumably by the spatial distribution of the islands, stochastic dispersals, and time since the formation of the barriers.

## INTRODUCTION

The interactions between historical and current processes that shape biogeographic patterns are a key component of biogeographical studies (e.g., Candolle, 1820; Good, 1974; Brown & Gibson, 1983; Cox & Moore, 1993; Crisci et al., 2003). In historical biogeography, biogeographic patterns are explained based on past processes that have persisted over very long periods of time, such as geological barriers formed by tectonic movements over millions of years (Crisci et al., 2003). However, current processes such as ongoing dispersals can affect or obscure evidence of historical patterns with migration across historical barriers (Wolf et al., 2001; Giller et al., 2004). Therefore, historical barriers and current dispersals compete in shaping biogeographic patterns; however, their relative importance is expected to vary depending on abiotic and biotic factors such as the spatial configuration of habitats and the dispersal ability of organisms. Increased knowledge of the shifting relative importance in relation to these factors would enhance our understanding of processes that structure biogeographic patterns. The Ryukyu Archipelago is a suitable study area for such studies.

The Ryukyu Archipelago is an assemblage of continental islands located between Kyushu Island of Japan and Taiwan (Fig. 1). This area is biogeographically interesting because several similar floristic differentiation patterns have been proposed, all of which are primarily explained by two historical barriers, the Tokara Tectonic Strait (Tokara Gap) and the Kerama Gap, which arose during the formation of the islands (Hara, 1959; Good, 1974; Maekawa, 1974; Shimabuku, 1984; Takhtajan, 1986; Kitamura et al., 1994). However, it is expected that plant groups whose seeds/fruits exhibit differing dispersal abilities display different floristic patterns because the effects of the historical barriers are dependent on dispersal ability. That is, a lower dispersal ability results in a greater amount of time that the historical effects are able to persist, whereas a higher dispersal ability results in fewer historical effects (Nekola & White, 1999). Therefore, it is thought that historical barriers can explain floristic differentiation patterns in plants with low dispersal ability, but not in plants with higher dispersal ability. In the latter case, it is expected that without an effective barrier to dispersal, the floristic differentiation would be correlated to the geographic distance between the islands, and that the floristic pattern is explained by this spatial effect.

**Fig. 1.**
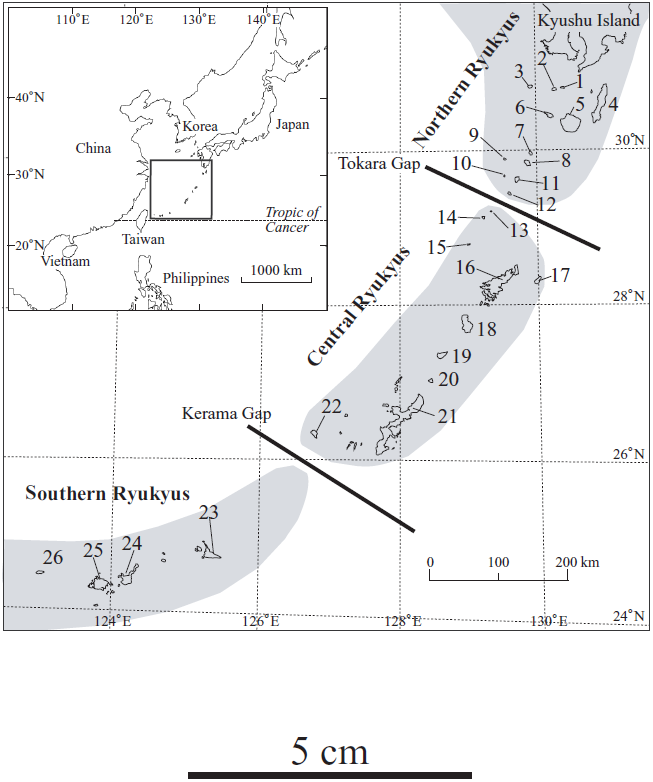
Location of the Ryukyu Archipelago in East Asia and distribution of the 26 islands analyzed in the archipelago. Segregation of the archipelago into the northern, central, and southern Ryukyus by the Tokara and Kerama Gaps is shown. Shaded areas indicate land configuration in the early Pleistocene after segmentation of the landbridge by the formation of the gaps, following Ota (1998). The names and the coordinates at the highest peaks (N, E) of the islands are as follows; 1, Takeshima (30° 48′, 130° 25′); 2, Ioujima (30° 47′, 130° 18′); 3, Kuroshima (30° 49′, 129° 56′); 4, Tanegashima (30° 38′, 130° 58′); 5, Yakushima (30° 20′, 130° 30′); 6, Kuchinoerabujima (30° 26′, 130° 13′); 7, Kuchinoshima (29° 58′, 129° 55′); 8, Nakanoshima (29° 51′, 129° 51′); 9, Gajajima (29° 54′, 129° 32′); 10, Tairajima (29° 41′, 129° 32′); 11, Suwanosejima (29° 38′, 129° 42′); 12, Akusekijima (29° 27′, 129° 35′); 13, Kodakarajima (29° 13′, 129° 19′); 14, Takarajima (29° 08′, 129° 12′); 15, Yokoatejima (28° 48′, 128° 59′); 16, Amamioshima (28° 17′, 129° 19′); 17, Kikaijima (28° 18′, 129° 58′); 18, Tokunoshima (27° 46′, 128° 58′); 19, Okinoerabujima (27° 22′, 128° 34′); 20, Yoronjima (27° 02′, 128° 25′); 21, Okinawajima (26° 43′, 128° 13′); 22, Kumejima (26° 22′, 126° 46′); 23, Miyakojima (24° 45′, 125° 22′); 24, Ishigakijima (24° 25′, 124° 11′); 25, Iriomotejima (24° 21′, 123° 53′); 26, Yonagunijima (24° 27′, 123° 00′).

To test this hypothesis, we compiled the distribution records of seed plants and grouped them into three dispersal-ability classes for each island of the Ryukyu Archipelago to examine the floristic patterns for each dispersal-ability class. We then used a multivariate approach to examine how the significance of the effects of the two historical barriers and the geographic distance differ with various dispersal-ability classes.

## MATERIAL AND METHODS

### Study area

The Ryukyu Archipelago is thought to have undergone extensive changes in land configuration during the Cenozoic, and there was more than one period of landbridge connection between the islands and surrounding landmasses (Kyushu to the north, and southeastern China via Taiwan to the south). This enhanced the range expansion of various lineages of terrestrial organisms in this region (Hatusima, 1975; Ota, 1998; Chiang & Schaal, 2006). It is proposed that by the Pliocene or early Pleistocene, the landbridge was segmented at two areas called the Tokara Gap and the Kerama Gap, where the sea is currently more than 1000 m deep (Kawana, 2002), as a result of tectonic movements and eustatic sea-level rise. It is thought that the islands have not since been connected across the gaps, even during the Quaternary glacial sea-level minima (Ota, 1998). Given this geohistory, many reports have demarcated the flora at one or both of the gaps (e.g., Hara, 1959; Good, 1974; Maekawa, 1974; Shimabuku, 1984; Takhtajan, 1986; Kitamura et al., 1994). For example, Takhtajan (1986) placed the demarcation line of the Japanese–Korean and Ryukyu provinces and that of the Ryukyu and Taiwanian provinces at the Tokara and Kerama Gaps, respectively. In addition, Good (1974) and Kitamura et al. (1994) regarded the Tokara Gap as the demarcation between the Sino–Japanese Region of the Boreal Kingdom and the Continental Southeast Asiatic Region of the Paleotropical Kingdom.

The spatial arrangement of the 26 islands examined in this study is shown in Fig. 1. These islands were selected because of the availability of reliable lists of seed plant flora. Based on the two gaps, the islands of the Ryukyu Archipelago are divided into three groups: the islands 1 to 12, 13 to 22, and 23 to 26 belong to the northern, central, and southern Ryukyus, respectively. The Ryukyu Archipelago is in the subtropics, or the transition from tropical to warm temperate zones (Good, 1974). The climate is moderate throughout the year with a mean temperature of approximately 15°C during the winter and 28°C in summer, and annual precipitation exceeds 2000 mm with no dry season. Therefore, the islands are covered in well-developed broad-leaved evergreen forests (Maekawa, 1974).

### Floristic data editing

The vascular plant flora of the Ryukyu Archipelago comprises approximately 1800 species (Hatusima & Nackejima, 1979); however, detailed records of each species for each island have not been compiled in one source. We referred to six literature sources for distribution records of native plants on the 26 islands (Hatusima & Amano, 1974, 1994; Hatusima et al., 1975; Niiro & Shinjo, 1989; Hatusima, 1991; Kawakubo & Tagawa, 1991). We compiled data sets for three dispersal-ability classes: high (269 wind-dispersed species with dust seeds and fruits having hairy pappi), low (200 gravity-dispersed species mostly with acorns and dehiscent fruits), and intermediate (224 bird-dispersed species mostly with sap-fruits). Each of the three classes included a similar number of species. Estimation of dispersal abilities was based on dispersal vectors estimated on the basis of seed/fruit morphology (e.g., Brown & Gibson, 1983). Infraspecific taxa were not separated to avoid taxonomic incongruence among the literature sources and to avoid giving intra- and inter-specific differences equal weighting.

For each dispersal-ability class, the pairwise floristic similarity between islands was calculated using Simpson’s similarity index (SI); SI = *γ β*^−1^, (*α* > *β*), where *γ* is the number of species shared between two islands, and *α* and *β* are the numbers of species on each of the two islands (Simpson, 1943). SI is one of the most frequently used similarity indices in biogeographic analyses (Brown & Gibson, 1983), and we selected this index because it does not give an improperly low similarity value when the areas (and hence the numbers of species) being compared differ greatly (Balgooy, 1971). Floristic dissimilarity distance was calculated by subtracting the value of SI from 1.

### Historical and spatial variables

The historical effect of the Tokara Gap was expressed as a categorical matrix using dummy variables; a value of 1 was given to matrix elements comparing two islands crossing over the Tokara Gap, and a value of 0 was given to matrix elements comparing two islands on either side of the Tokara Gap. The matrix of the Kerama Gap was constructed in the same manner. This is because the proposed dates of the formation of the gaps are very approximate (Ota, 1998), meaning that we cannot use these dates as accurate and reliable variables. Pairwise geographic distance (km) among the islands was calculated based on the geological coordinates at the highest points of each island (range = 11.3–1042.0 km; average distance = 353.2 km; SD = 284.0 km).

### Data analyses

To illustrate the floristic biogeographic pattern in each dispersal-ability class, clustering analyses of the islands were conducted based on floristic dissimilarity distances using the unweighted pair group method with arithmetic mean (UPGMA; Sneath & Sokal, 1973) with PAUP* 4.0b10 (Swofford, 2002).

To examine the significance of the effects of the Tokara and Kerama Gaps and the inter-island geographic distance on the variation in the floristic dissimilarity distance, we conducted partial Mantel tests. A Mantel test is used to test the statistical significance of correlations of distance matrices, where matrix elements are not independent, by randomization (Mantel, 1967; Smouse et al., 1986; Legendre et al., 1994). Partial Mantel tests are used to conduct partial linear regression analyses using a response matrix and two explanatory matrices to test the correlation between the response variable and one of the explanatory variables while controlling for the effect of the other explanatory variable to remove spurious correlation (Legendre, 2000; Bonnet & Peer, 2002; Tuomisto et al., 2003; Legendre et al., 2005; Lichstein, 2007). We conducted partial Mantel tests of the floristic dissimilarity distance with two pairs of explanatory variables, Tokara Gap and geographic distance, and Kerama Gap and geographic distance, using the software program ‘zt’ with 10000 randomizations (Bonnet & Peer, 2002).

To estimate the geographic extent of the significant correlation between the floristic dissimilarity distance and the geographic distance, we calculated a correlation coefficient *r* for given distance classes using GeneAlEx 6 software (Peakall & Smouse, 2006). Distance class sizes were determined to include a similar number of pairwise comparisons (32 or 33 island pairs) within each distance class. The calculated correlation coefficients were plotted as a function of geographic distance. Statistical significance of *r* was tested by 1000 randomizations for the null hypothesis of no correlation.

## RESULTS

### Floristic distance and clustering of islands

Mean values of the pairwise floristic dissimilarity distances were 0.240 (range = 0–0.576) in high, 0.187 (0–0.485) in intermediate, and 0.260 (0–0.641) in low dispersal-ability classes. UPGMA clusterings in three dispersal-ability classes are shown in Fig. 2. Seen in a broad perspective, the clustering patterns were similar and approximately congruent with the geographic arrangement of the islands in the archipelago. In all three classes, the islands of each of the northern, central, and southern Ryukyus were separated, with some exceptional small islands (islands 13–15, 26). Among the three regions, the islands of the central and southern Ryukyus were more closely connected to each other than to the islands of the northern Ryukyus.

**Fig. 2.**
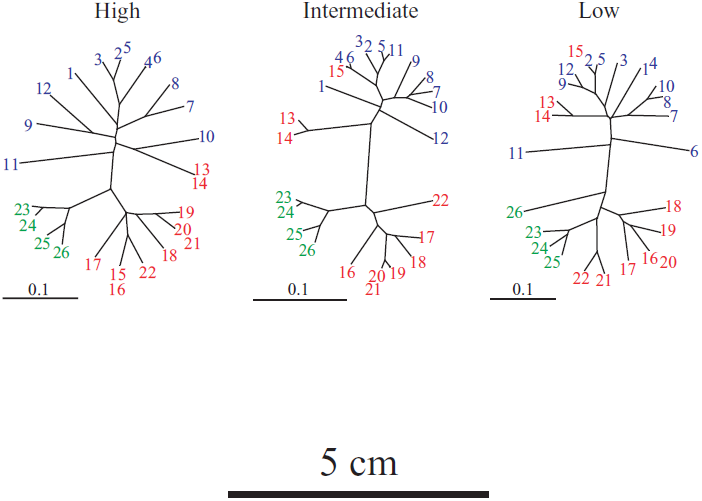
UPGMA clusterings of the 26 islands based on the floristic dissimilarity distance (1 – Simpson’s similarity index) in high, intermediate, and low dispersal-ability classes. Island numbers of the northern, central, and southern Ryukyus are colored blue, red and green, respectively. Island names are given in Fig. 1. The scale bars represent floristic dissimilarity distance of 0.1.

### Regression analyses

The results of the partial Mantel tests are shown in Table 1. In the high-dispersal ability class, the partial regression of the floristic distance matrix on the matrix of the Tokara Gap, controlling the effect of the geographic distance, was positive and significant (r = 0.31, p = 0.0001; in partial Mantel tests of this study, significance level 5% with Bonferroni adjustment was p = 0.05/12 = 0.0042), and so was the partial regression of the floristic distance matrix on the geographic distance, controlling the effect of the Tokara Gap (r = 0.31, p = 0.0002). The partial regression of the floristic distance matrix on the matrix of the Kerama Gap, controlling the effect of the geographic distance, was negative but not significant (r = -0.07, p = 0.1876), whereas the effect of the geographic distance, controlling the effect of the Kerama Gap, was significantly positive (r = 0.29, p = 0.0002). The results for the intermediate and low dispersal-ability classes were generally in accordance with one another. Specifically, the partial regression of the floristic distance matrix on the matrix of the Tokara Gap, controlling the effect of the geographic distance, was significantly positive (intermediate: r = 0.44, p = 0.0002; low: r = 0.18, p = 0.0037) and vice versa (intermediate: r = 0.38, p = 0.0001; low: r = 0.49, p = 0.0001). The partial regression of the floristic distance matrix on the matrix of the Kerama Gap, controlling the effect of the geographic distance, was significantly negative (intermediate: r = -0.28, p = 0.0003; low: r = -0.31, p = 0.0006) but the effect of the geographic distance, controlling the effect of the Kerama Gap, was significantly positive (intermediate: r = 0.49, p = 0.0001; low: r = 0.55, p = 0.0001).

**Table 1.**
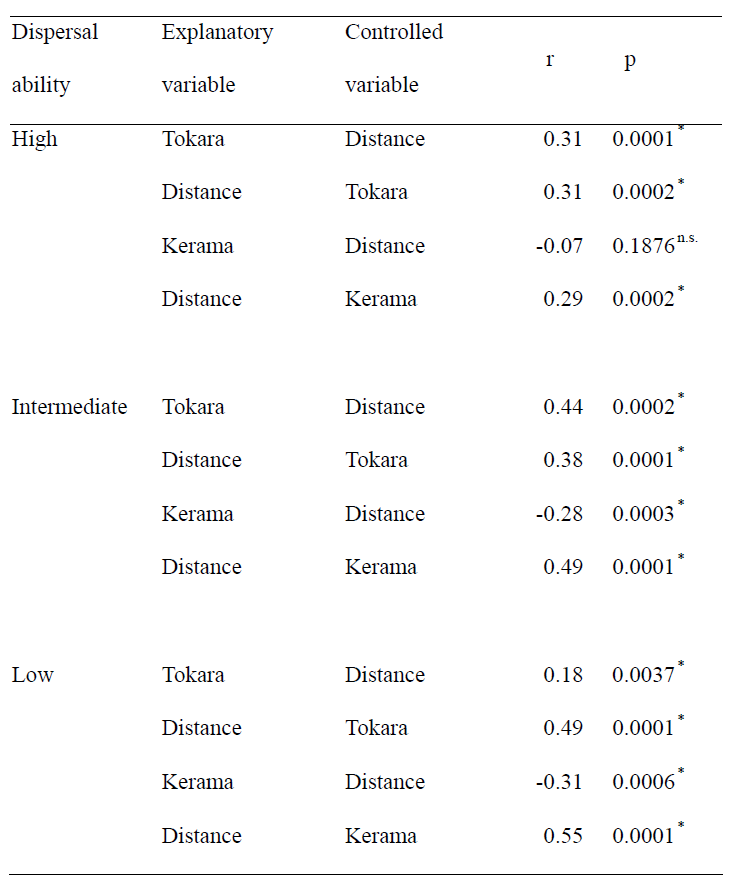
Summary of partial Mantel tests of floristic dissimilarity distance in three dispersal-ability classes. The tests were conducted with two pairs of explanatory variables, Tokara Gap-geographic distance, and Kerama Gap-geographic distance. The significance was tested with 10000 randomizations. Significance level 5% with Bonferroni adjustment for multiple comparisons was p = 0.05/12 = 0.0042. * = significant, n.s. = not significant

| Dispersal<br>ability | Explanatory<br>variable | Controlled<br>variable | r | p |
| --- | --- | --- | --- | --- |
| High | Tokara | Distance | 0.31 | 0.0001* |
|  | Distance | Tokara | 0.31 | 0.0002* |
|  | Kerama | Distance | -0.07 | 0.1876<sup>n.s.</sup> |
|  | Distance | Kerama | 0.29 | 0.0002* |
| Intermediate | Tokara | Distance | 0.44 | 0.0002* |
|  | Distance | Tokara | 0.38 | 0.0001* |
|  | Kerama | Distance | -0.28 | 0.0003* |
|  | Distance | Kerama | 0.49 | 0.0001* |
| Low | Tokara | Distance | 0.18 | 0.0037* |
|  | Distance | Tokara | 0.49 | 0.0001* |
|  | Kerama | Distance | -0.31 | 0.0006* |
|  | Distance | Kerama | 0.55 | 0.0001* |

### Geographic extent of spatial correlation

As shown in Figure 3, the analyses examining the geographic extent of the significant correlation between the floristic dissimilarity distance and the geographic distance generated very similar patterns in the three dispersal-ability classes. Specifically, the correlation coefficients were positive and significant in the first four distance classes (56, 104, 139, 186 km), with an x-intercept of approximately 250 km. The geographic extent of positive correlation can be influenced by the chosen distance class size (Double et al., 2005); however changing distance class sizes generated similar patterns (data not shown).

**Fig. 3.**
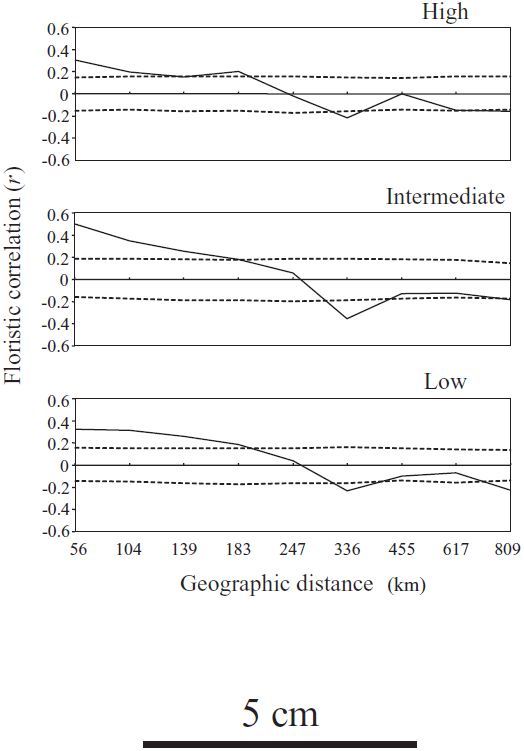
Plots of the floristic correlation coefficient (*r*) as a function of geographic distance in high, intermediate and low dispersal-ability classes. Each distance class includes the similar number of pairwise comparisons (32 or 33 island pairs). Dashed lines show the 95% confidence interval for the null hypothesis of no correlation, based on 1000 randomizations.

## DISCUSSION

### Dispersal ability and floristic patterns

The UPGMA analyses generated similar clustering patterns regardless of dispersal-ability class. This result rejects the hypothesis that plant groups with different dispersal abilities exhibit different floristic patterns. Biogeographic patterning results from each organism’s migration, which is a function of its dispersal ability and landscape resistance to the movement of the organism (Nekola & White, 1999). Therefore, the similar floristic patterns observed here are likely explained from these two viewpoints.

One explanation for our results is that the classification of dispersal ability was not adequate and there was no significant difference in the potential dispersal distances among the three classes. It is difficult to accurately estimate the dispersal ability of plants because evidence of dispersal, especially such a long-distance dispersal as to cross over sea, is not available for most plant species (Cain, 1974). Also, with respect to long distance dispersal, estimating dispersal abilities based on seed/fruit morphology might be misleading because the morphological adaptations of seeds/fruits that are typically used to identify the standard dispersal vector tend to govern short-distance dispersal but do not necessarily constitute the main mechanism responsible for long-distance dispersal (Nathan, 2006). Long-distance dispersal can occur by means of multiple stochastic dispersal vectors, even in the absence of specific adaptations for each vector (Bullock & Clarke, 2000; Myers et al., 2004). Conversely, some analyses examining the relationships between plant dispersal abilities and decrease-rate of floristic similarity with geographic distance have indirectly illustrated that plants with seeds/fruits adapted to bird-dispersal are more dispersal-limited than plants adapted to wind-dispersal (Nekola & White, 1999; Tuomisto et al., 2003). Therefore, classification of the plant dispersal abilities based on the seed/fruit morphology is not necessarily unreasonable. Assuming that our dispersal-ability classification was accurate, a second explanation for our results must consider the effect of landscape resistance.

Given that the landscape resistance is common to the analyses of the three dispersal-ability classes, it is likely responsible for the similar results in our analyses. Landscape resistance is caused by spatial configuration, particularly size and isolation of habitats. The more isolated and/or the smaller the habitat, the less effective dispersal will be (Nekola & White, 1999; Sklenář & Jørgensen, 1999). For example, in North American spruce-fir forests, isolated habitats restricted to the highest peaks, which are analogous to the habitats of islands, exhibit much stronger landscape resistance than more contiguous habitats (Nekola & White, 1999). In the Ryukyu Archipelago, it is very plausible that the isolated distribution of relatively small islands is causing strong landscape resistance. As shown in Fig. 3, the distance at which the correlation between floristic dissimilarity distance and geographic distance becomes effectively zero was approximately 200 km, regardless of the dispersal-ability class. This means that the geographic extent of the correlation is confined to a comparatively small area while the archipelago extends for more than 1300 km. We propose that because the landscape resistance is so strong that migration of plants is severely restricted regardless of their dispersal abilities, the plant groups whose potential dispersal distances in a contiguous landscape are different generate similar differentiation patterns.

### Significance/insignificance of historical effects

The partial Mantel tests indicated that the floristic differentiations among islands were explained by both historical effect and geographic distance in all three dispersal-ability classes. This result rejects the hypothesis that historical barriers can not explain floristic differentiation patterns in plants with higher dispersal ability. The UPGMA analyses indicated that the northern, central, and southern Ryukyus were essentially separated. This separation pattern seems to be consistent with the hypotheses of the preceding studies, which explain the floristic differentiation pattern primarily by the historical gaps (Hara, 1959; Good, 1974; Maekawa, 1974; Shimabuku, 1984; Takhtajan, 1986; Kitamura et al., 1994). However, our partial Mantel tests indicated that geographic distance and the Tokara Gap have exhibited significantly positive effects but that the effects of the Kerama Gap were insignificant or significantly negative. The significantly positive effect of the Kerama Gap was spurious. Note that the geographic distances between islands across the Kerama Gap are much larger than those of the Tokara Gap (the shortest distances across each gap are 227.6 km between islands 22 and 23 and 36.6 km between islands 12 and 13; Fig. 1). However, the UPGMA analyses indicated that the islands of the central and southern Ryukyus were more closely connected to each other than to the islands of the northern Ryukyus. Therefore, the explanation for the insiginificant effect of the Kerama Gap in high dispersal-ability class is that the floristic dissimilarity distance across the Kerama Gap was no more that what was explained by the effect of the accompanying large geographic distance. Similarly, the explanation for the negative effects in the intermediate and low dispersal-ability classes is that the floristic dissimilarity distance across the gap was less than what was expected from the large geographic distance, for which the effect of the Kerama Gap was adjusted to be negative. This is because that although the significantly positive effect of the Kerama Gap was spurious, the negative effects in intermediate and low dispersal-ability classes could also be spurious because it is unlikely that the formation of the Kerama Gap increased the floristic similarity among the islands across the gap, considering that the landbridge had played a significant role in the migration of the plants (Hatusima, 1975; Chiang & Schaal, 2006). Thus, the effect of the Kerama Gap would actually be positive but indistinguishable compared to the effect of the large geographic distance.

When we consider that the Tokara Gap is a very narrow strait (the two closest islands across the gap are only 36.6 km apart today), the significant effect of the Tokara Gap requires an explanation as to why ongoing inter-island dispersal has not completely obscured the effect of the past formation of the gap. The duration of the effects of a historical barrier is determined by the amount of time required for organisms to cross it (Nekola & White, 1999). The Tokara Gap is thought to have formed during the Miocene or at the latest in the early Pleistocene, roughly more than two million years ago (Ota, 1998). A recent quantitatively derived understanding of the long-distance dispersal of plants may help us understand the persistence of the historical effects of the Tokara Gap. The expected time for an effective dispersal, i.e., the successful establishment of one reproductive individual is calculated as the inverse of the product of seed arrival probability and seed-to-adult survival probability (Clark et al., 1999). For example, with a hypothetical source population (10^6^ individuals, each with an annual fecundity of 10^4^ seeds and with mean dispersal distance of 50 m), the expected time for a single effective dispersal event to occur beyond 150 km is longer than 100 billion years under the mean trend. Conversely, an effective long-distance dispersal event beyond 415 km, expected to occur once in almost 10^13^ years under the mean trend, may occur once in 10 years as a result of events that break the rules, such as hurricanes, tornadoes, and tropical cyclones. Similarly, an effective long-distance dispersal event beyond 34 km may occur once in almost 10^8^ years under the mean trend but once a year under exceptional conditions (Clark et al., 1999). Long-distance dispersal is an extremely rare phenomenon even at the scale of a few tens of km or a few hundred km, although nonstandard mechanisms may help realize it (Nathan, 2006). Although these are rough estimates for certain types of plants, they still serve as useful references. The time required for long-distance dispersal to cross the Tokara Gap under the mean trend would be much longer than the time since the formation of the gap (> 2 million years), although this time would have allowed some species to cross the gap with nonstandard mechanisms such as summer typhoons. Thus, it is likely that the historical effects of the Tokara Gap have been preserved because of the rarity of long-distance dispersal across the gap beyond 36.6 km under the mean trend with the time being limited. Therefore, the significance of the historical barriers is not determined by the plant dispersal abilities but presumably by the spatial distribution of the islands, stochastic dispersals, and time since the gap formation.

## ACKNOWLEDGEMENTS

We thank Drs. A. J. Davis (Max Planck Institute for Chemical Ecology), N. Kawakubo (Gifu University), S. Matsumura (University of the Ryukyus) and H. Ota (University of the Ryukyus) for their valuable comments. This study was supported by a Research Fellowship for Young Scientists from the Japan Society for the Promotion of Science (JSPS) to K.N., a Grant-in-Aid for Scientific Research (C) (no. 17570083) to M.Y., and a grant for the 21st Century COE program of the University of the Ryukyus.

